# Bacterial ecology and evolution converge on seasonal and decadal scales

**DOI:** 10.1101/2024.02.06.579087

**Authors:** Robin R. Rohwer, Mark Kirkpatrick, Sarahi L. Garcia, Matthew Kellom, Katherine D. McMahon, Brett J. Baker

## Abstract

Ecology and evolution are often viewed as distinct processes, which interact on contemporary time scales in microbiomes. To observe these processes in a natural system, we collected a two-decade, 471-sample freshwater lake time series, creating the longest metagenome dataset to date. Among 2,855 species-representative genomes, diverse species and strains followed cyclical seasonal patterns, and one in five species experienced decadal shifts in strain composition. The most globally abundant freshwater bacterium had constant species-level abundance, but environmental extremes appeared to trigger a shift in strain composition and positive selection of amino acid and nucleic acid metabolism genes. These genes identify organic nitrogen compounds as potential drivers of freshwater responses to global change. Seasonal and long-term strain dynamics could be regarded as ecological processes or equivalently as evolutionary change. Rather than as distinct processes that interact, we propose a conceptualization where ecology and evolution converge along a continuum to better describe change in diverse microbial communities.

## Main Text

Microbial communities allow us to observe eco-evolutionary dynamics in real-time due to the short lifespans and large population sizes of microbes. Real-time evolution was famously observed in the *E. coli* long-term evolution experiment^1^, but no similar long-term observations exist for natural, ecologically complex systems. Here we introduce a two-decade, 471-sample microbial time series from a freshwater lake, the TYMEFLIES dataset, which allows us to directly observe ecology and contemporary evolution in a natural ecosystem. By reconstructing tens of thousands of metagenome-assembled genomes (MAGs), we found that ecology and evolution both unfold at short, seasonal time scales as well as longer-term decadal time scales. In genomes from the most abundant freshwater bacterium, *Nanopelagicaceae*, evolutionary change coincided with environmental extremes. While species-level abundance remained constant, strain composition shifted coincident with an increase in genes under positive selection. Research on eco-evolutionary dynamics focuses on feedbacks between distinct processes of ecology and evolution^2–4^. In our microbial data, however, these processes were difficult to distinguish; ecological dynamics appeared to occur between strains, but the strains themselves were inferred from observations of genomic change. Consistent with the ambiguity of the microbial species concept^5^, our observations suggest that it is not possible to cleanly delineate between ecological and evolutionary processes in microbial communities. Therefore, we propose an adjusted conceptualization, where ecology and evolution converge along a continuum.

## The TYMEFLIES dataset

We collected 471 samples over 20 years from Lake Mendota (WI, USA)^6^ and obtained shotgun DNA libraries (**Fig. 1A**, Supplementary Data 1). We refer to these Twenty Years of Metagenomes Exploring Freshwater Lake Interannual Eco/evo Shifts as the TYMEFLIES dataset. By cross-mapping reads from ∼50 metagenomes to each single-sample metagenome assembly, we obtained a total of 85,684 genome bins, 30,389 of which were medium or high quality (> 50% completeness and < 10% contamination)^7^. We clustered these 30,389 bins at 96% average nucleotide identity (ANI) and obtained 2,855 clusters from which we chose representative MAGs^8^ (Supplementary Data 2). Several previous studies have found an emergent species boundary at similar ANI cutoffs^9–11^, and we observed a rapid increase in the number of clusters above the 96% ANI cutoff. In this study, we treat the representative MAGs from each 96% ANI cluster as bacterial species and refer to sub-species delineations identified in the mapped metagenomic reads as strains^12^.

**Fig. 1.**
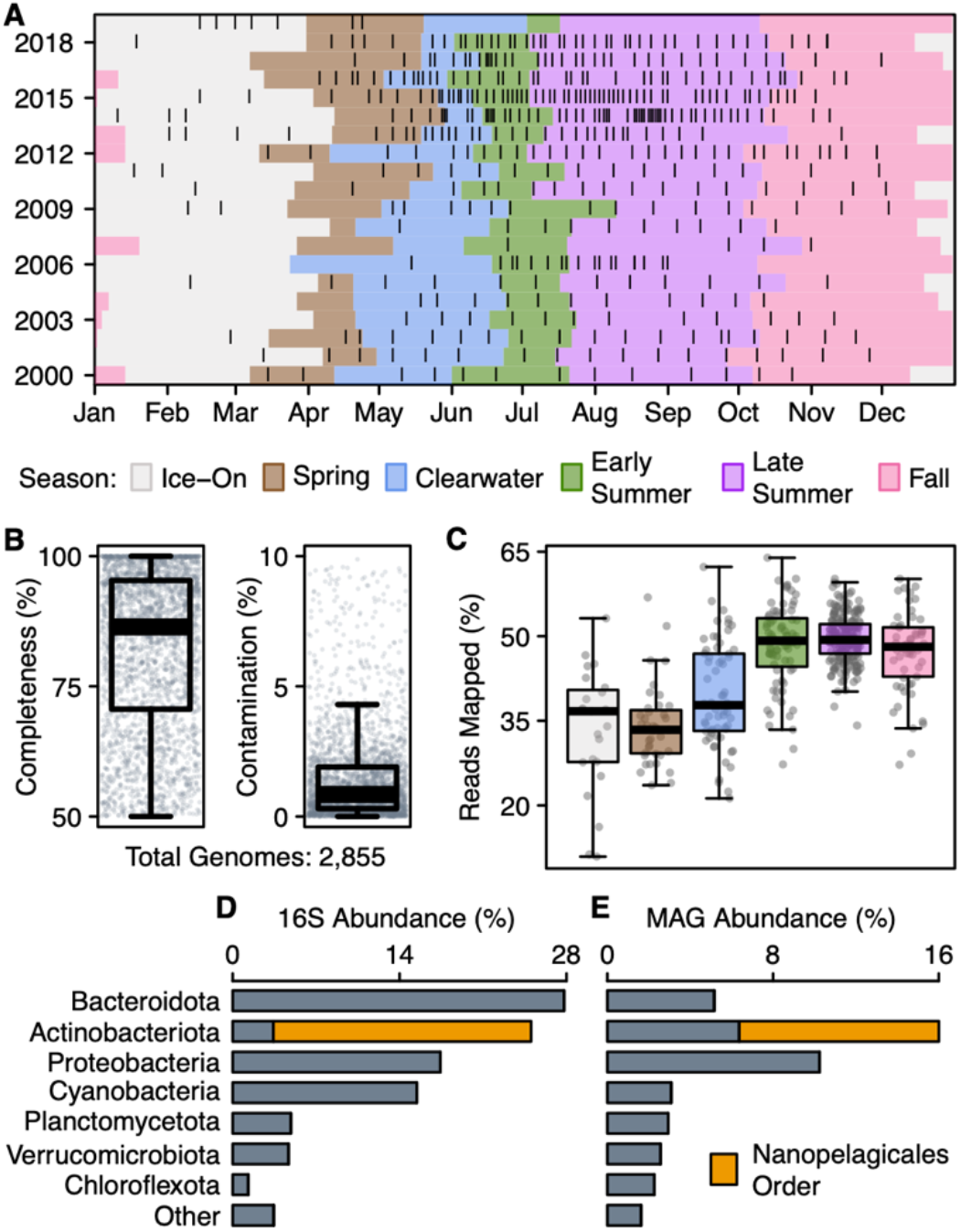
The TYMEFLIES dataset. **A)** Metagenome sample dates are indicated by black vertical lines. Microbial seasons^22^ are indicated by colored shading. **B)** Quality of the 2,855 representative genomes obtained after clustering to 96% ANI. We treat these genomes as species. **C)** Percent of metagenome reads from each sample that mapped to all reference genomes with an ANI ≥ 93%. Samples are grouped by season to highlight how well the reference genomes reflect each seasonal community. **D)** Rank abundance of phyla as measured by 16S rRNA gene amplicon sequencing^6^. The abundant *Nanopelagicales* order of Actinobacteria is highlighted. **E)** Abundance of phyla in the TYMEFLIES reference genomes, quantified as the mean relative abundance normalized by genome size and sequencing depth. The *Nanopelagicales* order is again highlighted.

The representative MAGs have high estimated completeness (median 86%) and low contamination (median 0.9%) (**Fig. 1B**), and reflect the abundant members of the lake’s bacterial community, especially in well-sampled seasons (**Fig. 1C**). Using a 16S rRNA gene amplicon dataset from the same timeseries^6^ as a reference for the expected community composition (**Fig. 1D**), we found that our representative MAGs comprise most of the abundant taxa (**Fig. 1E**). Moreover, we obtained 168 representative MAGs from the *Nanopelagicales* order, which is the most abundant order in Lake Mendota and accounts for 22% of the amplicon reads and 10% of the mapped metagenomic reads. Similar to SAR11 bacteria in the oceans, this freshwater lineage is abundant in lakes globally^13^, difficult to culture^14^, and typically has highly streamlined genomes^15^.

### Seasonal ecology and evolution

Lake Mendota has been the focus of limnological research since the late 1800s and has been routinely sampled since 1984 by the North Temperate Lakes Long-Term Ecological Research program (NTL-LTER)^16^. Microbial sampling began in 2000 as part of an NSF microbial observatory^17^. From this long history of research, we know the lake follows a consistent annual phenology, and that phenological patterns are changing in response to climate change and invasive species^18–21^. Rohwer *et. al*^22^ found that these phenological dynamics extend to the bacterial community. To confirm that phenological abundance patterns also exist in our more finely resolved bacterial species, we identified annual peaks in species relative abundance using periodograms (magnitude of Fourier transforms). After limiting this temporal analysis to the subset of 1,474 species that occurred at least 30 times over at least 10 years, we found that 72% of them have consistent seasonal abundance patterns (**Fig. 2A**).

**Fig. 2.**
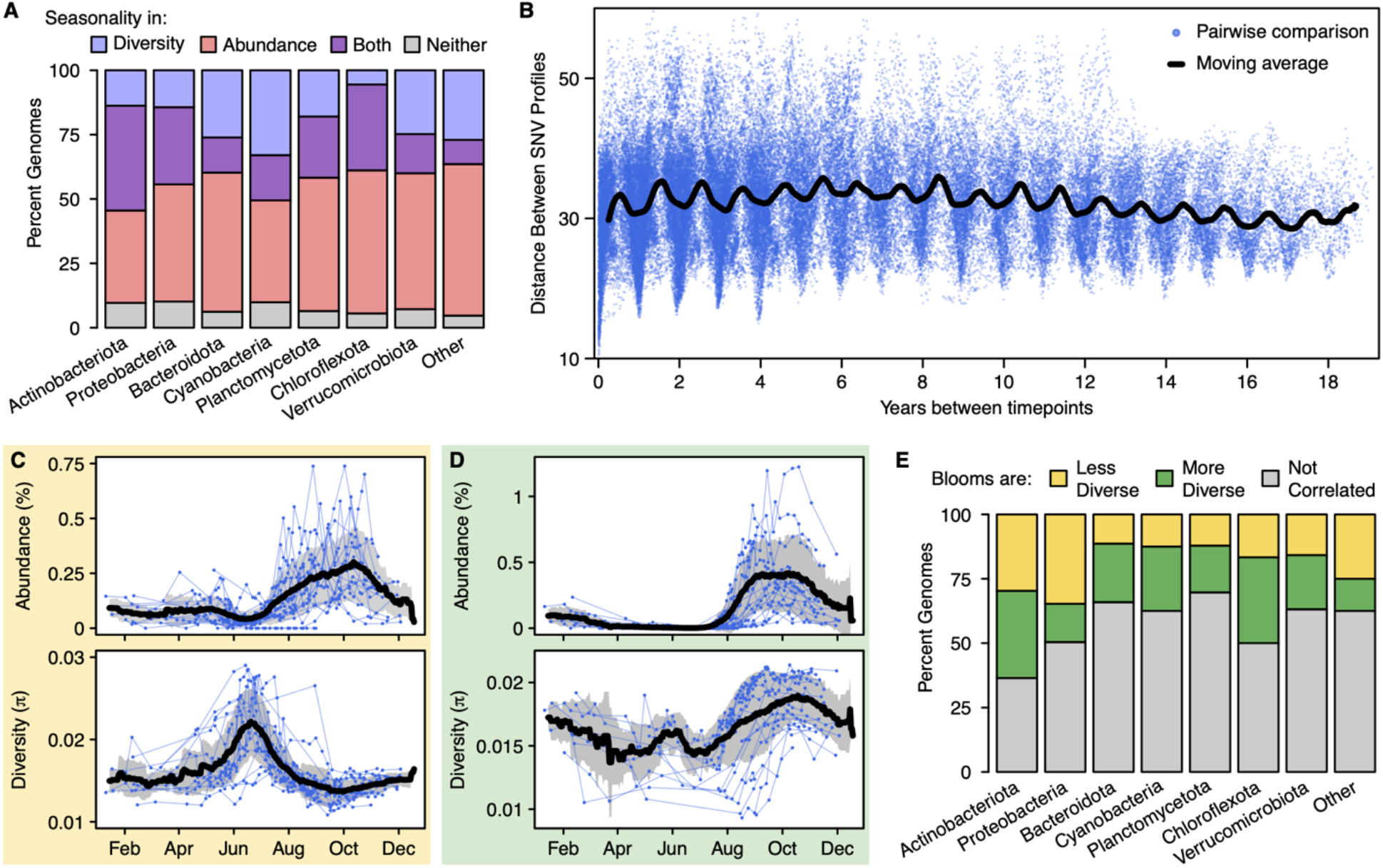
Bacterial seasonality at the sub-species level. **A)** The percent of species with seasonality in nucleotide diversity and abundance (a centered log ratio transform was applied to relative abundances). The 1,474 reference species that occurred at least 30 times were included in this analysis. **B)** A time decay plot of the Euclidean distances between the SNV profiles of an abundant species in the *Nanopelagicus* genus (ME2017-06-13_3300043469_group7_bin14). A smaller distance between SNV profiles indicates that the strain composition is more similar. Each blue point represents a pairwise comparison between two sample dates, with the time between those dates on the x-axis. The black line is a 6-month moving average, drawn to highlight the annual periodicity of strain similarities. **C)** An example of a less diverse bloom, where nucleotide diversity decreases as relative abundance increases. Displayed is an abundant species in the *Planktophila* genus (ME2011-09-04_3300044729_group3_bin142). **D)** An example of a more diverse bloom, where nucleotide diversity increases as abundance increases. Displayed is an abundant species (ME2012-08-31_3300044613_group4_bin150) in the *Nanopelagicaceae* family, MAG-120802 genus. **E)** The distribution of bloom diversity patterns across the 365 species that had seasonality in both abundance and nucleotide diversity.

To determine whether evolutionary dynamics (*i*.*e*. changes in allele frequency within the species) also unfold seasonally, we mapped reads from each sample against each species’ reference genome and identified shifts in strain composition from changes in nucleotide diversity (π) and allele frequencies at single nucleotide variants (SNVs). We found that 33% of the 1,474 species displayed consistent seasonal nucleotide diversity patterns (**Fig. 2A**). To gain greater resolution of the strain composition of the 236 species abundant enough over time to reliably call SNVs (median coverage > 10x), we created a “SNV profile” for each date with the frequencies of the reference alleles. For each species, we calculated the Euclidean distance between every date’s SNV profile (**Fig. 2B**). We found that 80% of these 236 abundant species had consistent phenological patterns in their strain composition. This demonstrates that phenological patterns evident in the bacterial community extend to the finest possible taxonomic resolution. Several short-term freshwater studies have also observed changes in strain composition on seasonal timescales^23,24^. Phenological patterns in sub-species strains similar to those at the species-level suggest ecological processes may shape bacterial strain composition, but these changes are evidenced by intraspecific genomic change and could thus also be interpreted as seasonal evolution.

Given the ubiquity of seasonal patterns in both species abundance and sub-species diversity, we asked if they were correlated. We quantified whether a species’ “bloom” in abundance consisted of fewer strains or more strains than its baseline composition. Of the 365 species with seasonal patterns in both abundance and nucleotide diversity (purple bars in **Fig. 2A**), we found both scenarios were common; 21% of these species had less diverse blooms (**Fig. 2C** and yellow bars in **Fig. 2E**), while 19% had more diverse blooms (**Fig. 2D** and green bars in **Fig. 2E**). Further, all abundant phyla demonstrated an even mix of both bloom types (**Fig. 2E**). A lower-diversity bloom suggests that a subset of strains outcompeted the others, while a higher-diversity bloom suggests that micro-niches allowed rarer strains to gain abundance, resulting in higher strain diversity^25^ due to a more even strain composition. This is in agreement with a previous study that found both overlapping and distinct niches within freshwater bacterial species^26^. The prevalence of both bloom diversity patterns suggests ecological processes drive changes in allele frequencies.

### Long-term ecology and evolution

Long-term changes can be masked by seasonal oscillations, lost in what is referred to as the “invisible present”^27^. The unprecedented length of the TYMEFLIES metagenome dataset provides a unique lens into the invisible present, enabling the identification of overlayed long-term patterns. To find long-term changes in strain composition, we developed a classifier trained on the distance between each date’s SNV profile and the SNV profile of that species’ first occurrence in the timeseries. We trained this classifier on 11 examples of manually identified temporal patterns, and then applied it to all 263 species with sufficient abundance to reliably call SNVs. Our classifier identified gradual change (**Fig. 3A**), which may arise from genetic drift or in response to a slow press disturbance. It also identified abrupt change (**Fig. 3B** and **C**), which may arise in response to a new stable state after a tipping point, or from a sudden environmental shift^28,29^. Among instances of abrupt change, we identified step changes (**Fig. 3B**), where the new strain composition persisted during the remainder of our time frame, as well as patterns of disturbance with resilience (**Fig. 3C**), where the strain composition recovered to baseline.

**Fig. 3.**
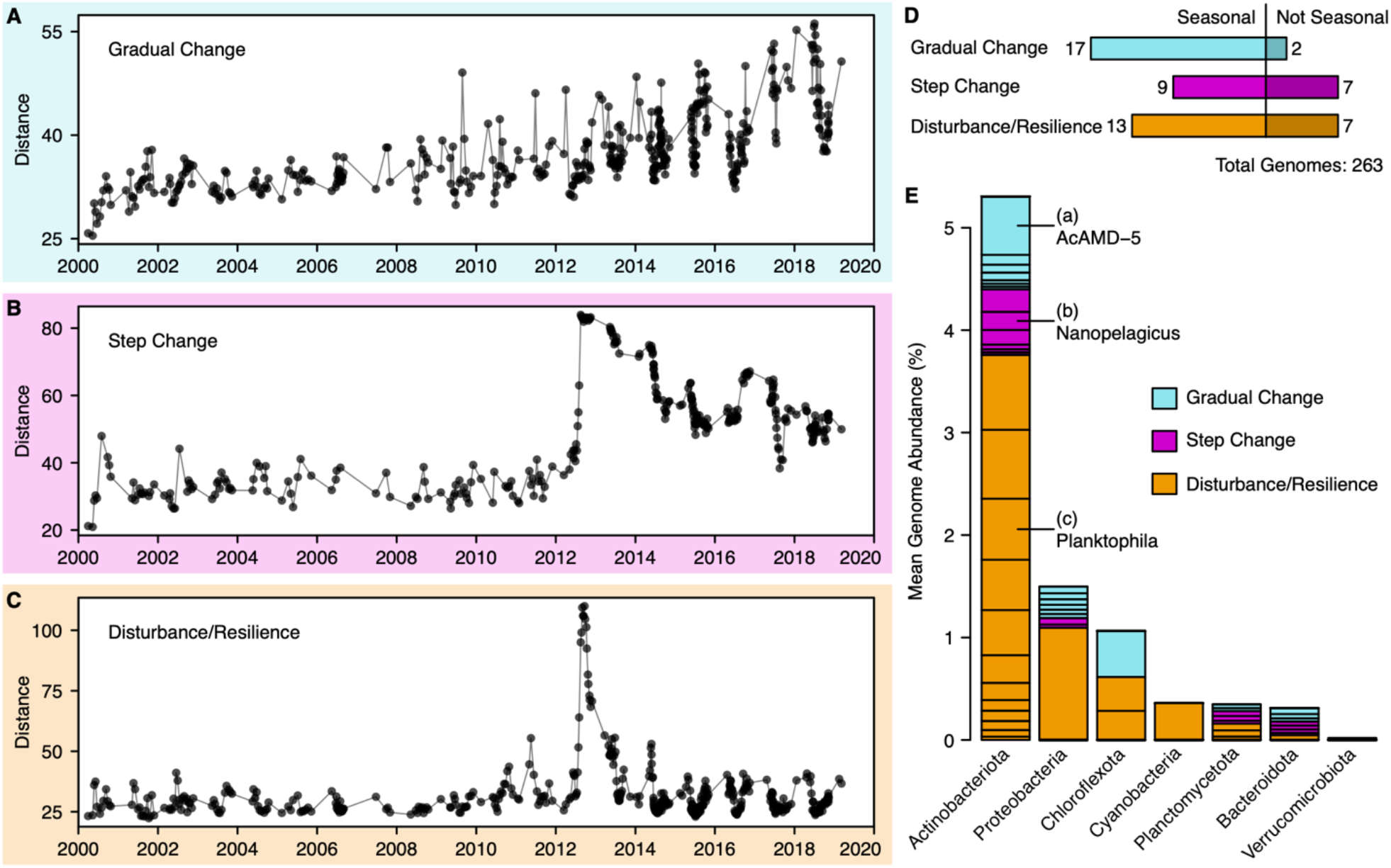
Long-term changes in strain composition. **A)** An example of long-term, gradual change in strain composition. Points indicate sample dates, and distance refers to the Euclidean distance between a species’ SNV profile on that sample date and its first occurrence in the time series. A species in the *Nanopelagicales* order, AcAMD-5 family is shown (ME2005-06-22_3300042363_group2_bin84). **B)** An example of an abrupt step change in strain composition in a species in the *Nanopelagicus* genus (ME2011-09-21_3300043464_group3_bin69). **C)** An example of a disturbance/resilience pattern, where an abrupt change in strain composition is followed by recovery to the original strain composition, in a species in the *Planktophila* genus (ME2015-07-03_3300042555_group6_bin161). **D)** Long-term change patterns often overlayed seasonal patterns. Of the 263 species abundant enough to observe their SNV profiles, 39 had both long-term and seasonal patterns while 16 had only long-term patterns. **E)** The distribution of long-term patterns across phyla. Each species that underwent long-term change is indicated by a section of the phyla’s bar, scaled by the mean abundance of that species. The sections corresponding to the examples highlighted in **A-C** are labelled.

We found that 21% of the most abundant species experienced one kind of long-term change in their SNV profiles during our 20-year study period, and these changes overlayed both seasonal and acyclical short-term dynamics (**Fig. 3D**). Abrupt change was almost twice as common as gradual change (seen in 36 vs. 19 species), and resilience was only slightly more common than a lasting step change (20 vs. 16 species) (**Fig. 3D**). The three long-term change patterns were found in many abundant species distributed across phyla (**Fig. 3E**). Many species in the Actinobacteriota phylum were abundant enough to include in this analysis, providing a detailed view of change in these common freshwater heterotrophs. Long-term changes in SNV profiles reflect shifts in intraspecific strain composition, which is typically attributed to evolutionary processes^30^. The fact that during our observation period over a fifth of the species experienced long-term changes in their SNV profiles emphasizes the importance of including contemporary evolutionary change in our understanding of microbial ecology.

### Abrupt changes in *Nanopelagicaceae*

In general, related species did not change in unison with each other, suggesting that the drivers of evolutionary change are highly specific (**Fig. 4A**). One exception is an abrupt change event that impacted seven species within the *Nanopelagicaceae* family (acI) in 2012, specifically species in the *Nanopelagicus* and *Planktophila* genera (acI-B and acI-A). This is the most abundant family in Lake Mendota and in freshwaters globally^13^, and the 127 *Nanopelagicaceae* species we recovered accounted together for 8% relative abundance on average. Five of these *Nanopelagicaceae* species displayed resilience to the abrupt change, while two experienced lasting step changes in strain composition.

**Fig. 4.**
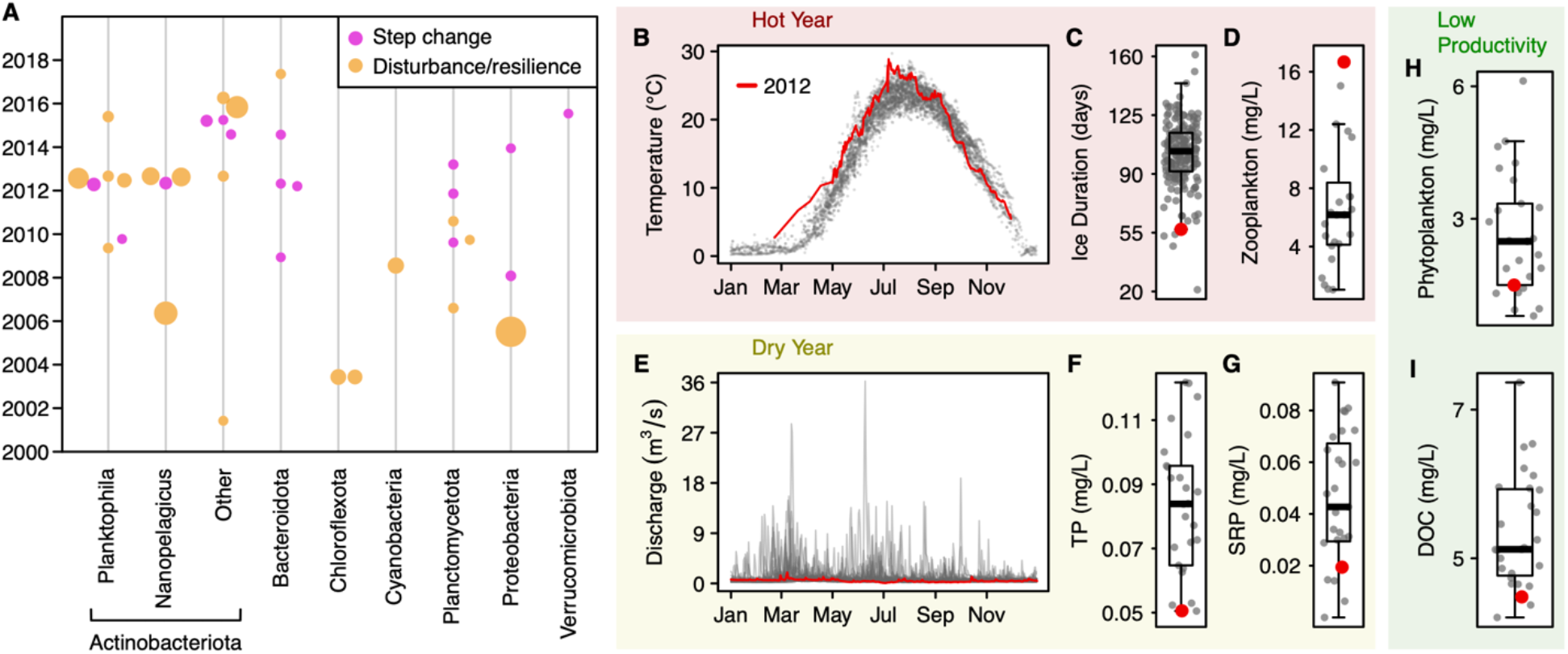
Abrupt changes in *Nanopelagicaceae* strain composition coincide with environmental extremes in 2012. **A)** Dates of all abrupt changes in strain composition arranged by phyla. Most changes were isolated events, but multiple species from two abundant genera of Actinobacteriota, *Planktophila* and *Nanopelagicus*, experienced abrupt change in 2012. Point size is scaled by species abundance. **B)** Unusually high epilimnion water temperatures during spring and summer 2012 (relative to 1894 – 2019). **C)** The preceding winter had an unusually short ice duration (relative to 1853 – 2023). **D)** Total zooplankton biomass (excluding predatory *Bythotrophes* and *Leptodora*) was unusually high, likely enabled by warm early spring temperatures (relative to 1995 – 2018). **E)** Discharge from the Yahara River, the main tributary to Lake Mendota, was unusually low and lacked high runoff events typical after storms and spring snowmelt (relative to 1989 - 2021). **F)** Total phosphorus, and **G)** soluble reactive phosphorus were low (relative to 1995 – 2021), likely due to low sediment transport. **H)** Low phytoplankton biomass (relative to 1995 – 2020), likely resulting from both high zooplankton grazing and low nutrient availability. **I)** Low dissolved organic carbon (relative to 1996 – 2022), likely a result of low phytoplankton abundance.

A myriad of possible environmental variables could have driven this event. A leading candidate is extreme weather, as Lake Mendota was unusually warm and dry in 2012. The lake experienced high epilimnion water temperatures during spring and summer, with the hottest July on record since 1894^22^ (**Fig. 4B**), the fifth shortest winter ice duration on record since 1856^31^ (**Fig. 4C**), the eighth lowest annual discharge from its major tributary on record since 1976 and the second lowest peak discharge^32^ (**Fig. 4E**). These environmental conditions led to top-down and bottom-up controls on the lake’s primary productivity. The highest spring zooplankton abundance since measurements began in 1994^33^ (**Fig. 4D**) was likely a result of the mild winter and spring^34^ which allowed zooplankton, including the prolific grazer *Daphnia pulicaria*, to establish early. Low total phosphorus and soluble reactive phosphorus (**Fig. 4F**-**G**) was likely a result of low external nutrient loading associated with mild discharge events^35^. The resulting combination of high zooplankton grazing and low phosphorus, typically the limiting nutrient in lakes, may be responsible for low phytoplankton biomass (**Fig. 4H**), which in Lake Mendota is dominated by Cyanobacteria during summer^36^. Lake Mendota’s dissolved organic carbon (DOC) is primarily provided by phytoplankton^37^, consequently DOC was also low in 2012 (**Fig. 4I**). Lake heatwaves are predicted to become hotter and longer with climate change^38^, and these observations suggest that the intense epilimnetic heat waves during 2012 had cascading effects on lake biogeochemistry that extended to the level of bacterial strains.

Another possible driver is the irruption of the invasive zooplankton spiny water flea (*Bythorephes cedertrömii*) in 2009, which itself was driven by an unusually cool summer^39^. This major disturbance resulted in a trophic cascade that decreased water clarity^21,40^, increased lake anoxia^33^, and shifted the bacterial community composition^22^. Although the abrupt changes in strain composition of seven *Nanopelagicaceae* species were not observed until three years later, lag effects are common in complex ecosystems^41^. In contrast to the 2009 species invasion, we did not see bacterial community-level shifts corresponding to the 2012 extreme weather, but environmental drivers of strain dynamics may be highly specific. Ecosystem-wide drivers like these two disturbances can have cascading and interacting effects on nutrient and carbon dynamics, which in turn impact bacteria. The observed long-term intraspecific changes suggest that such ecological drivers are also drivers of evolutionary change, further emphasizing how ecology and evolution are intertwined.

### Evolutionary signals in a *Nanopelagicus*

To understand the dynamics of abrupt evolutionary change, we further examined one of the abundant species, a *Nanopelagicus* (acI-B), that experienced a step change in strain composition in August 2012 (**Fig. 3B**). An NMDS ordination of its SNV profiles indicated the strain composition changed abruptly at that time and settled into a new composition after a period of adjustment in 2012 and 2013 (**Fig. 5A**).

**Fig. 5.**
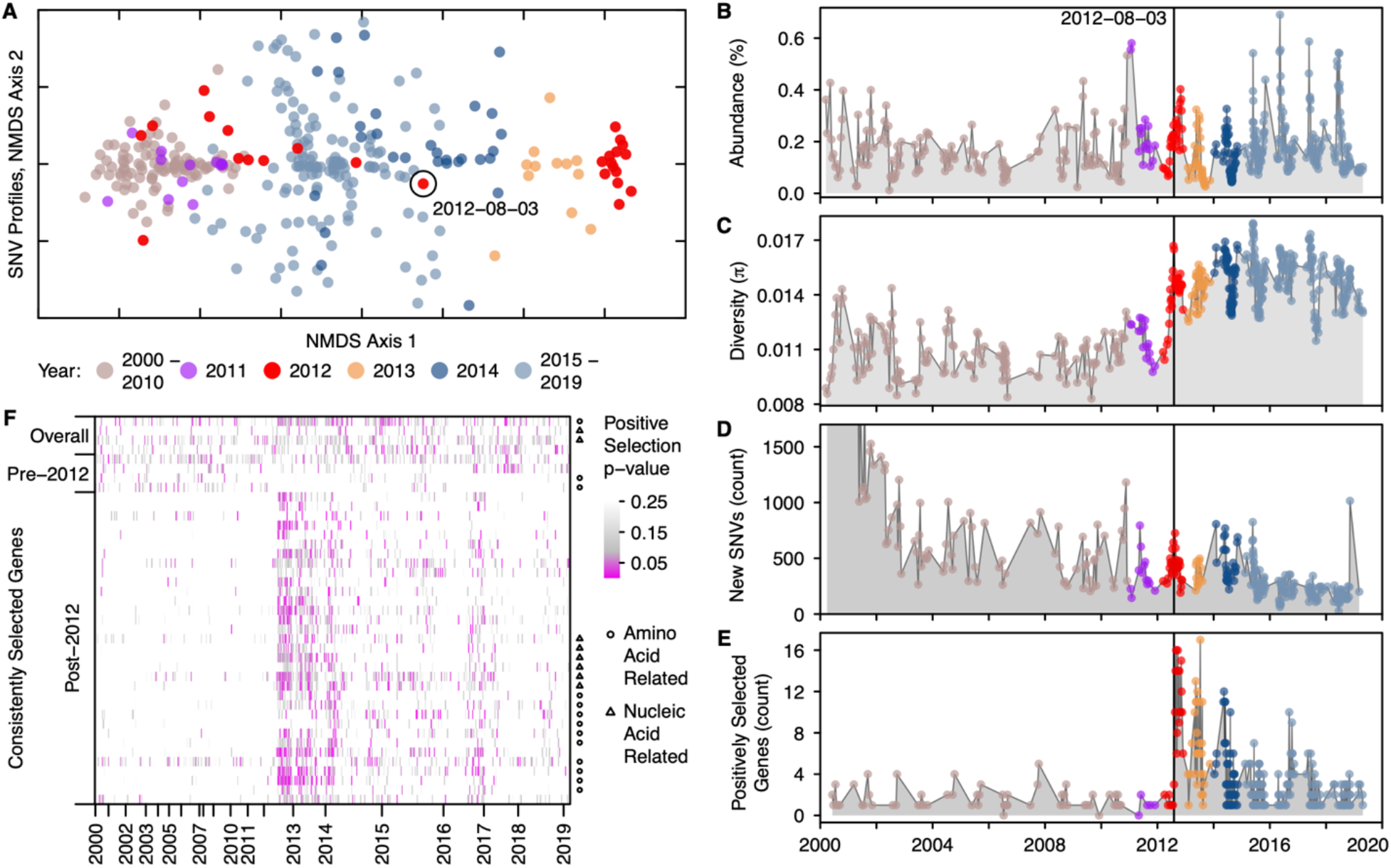
Step change in strain composition coincides with more genes under selection. **A)** An abundant *Nanopelagicus* species experienced a step change in strain composition in 2012 (ME2011-09-21_3300043464_group3_bin69, see also **Fig. 3B**). Samples with more similar SNV profiles appear closer on this NMDS plot. Years 2000-2011 cluster together and are distinct from years 2014-2019, which cluster separately. A sudden change in strain composition occurred on August 3, 2012. **B)** Despite the abrupt change in strain composition, the relative abundance of this species remained constant over time. **C)** Concurrent with the shift in strain composition, nucleotide diversity increased and then remained high, indicating that the new equilibrium was comprised of a more diverse assemblage of strains. **D)** The absence of a spike in the number of new SNVs suggests that an increase in the evenness of existing strains occurred, rather than the introduction of new strains. **E)** Concurrent with the shift in strain composition, the number of genes under positive selection also increased (McDonald-Kreitman F-statistic p-value < 0.05). **F)** Occurrence of consistently selected genes in all the samples, in the pre-2012 period, and in the post-2012 period. X-axis indicates samples over time and Y-axis indicates genes. Shading indicates the significance level of positive selection. Amino acid-related genes and nucleic acid-related genes are indicated on the right axis. Full annotations are available in Supplementary Data 3. Note that the X-axis is evenly spaced by sample, so that years with more samples take up more space.

The relative abundance of this species was quite constant throughout our 20-year observation period (**Fig. 5B**), typically with higher abundances during the spring clearwater phase. The step change in strain composition (**Fig. 3B**) coincided with one in genome-wide nucleotide diversity (**Fig. 5C**). These patterns could result from the introduction of a new strain or with an increase in the evenness of existing strain abundances. To distinguish between these hypotheses, we counted the number of previously unobserved SNVs in the mapped reads of every sample. We did not see large spikes in new SNVs in 2012 (**Fig. 5D**), suggesting that the step change reflects shifts in the relative abundances of existing strains.

This interpretation is consistent with a dramatic increase in the number of genes under positive selection that occurred at this time (**Fig. 5E**). As the relative abundances of some strains increase, alleles specific to them appear to undergo partial (or “soft”) selective sweeps. If strain composition re-equilibrated, this signal would die out. However, the increase in the number of genes under selection persisted (**Fig. 5E**). This could arise from continuing fluctuations in strain abundances, consistent with the larger distances between SNV profiles seen after the step change (**Fig. 5A**). To identify candidate loci that reflect the phenotypic differences between strains driving adaptations, we sought genes that consistently showed signs of being positively selected over the entire timeseries, only during the pre-2012 period, and only during the post-2012 period. Four genes were consistently selected both pre- and post-2012, four genes were consistently selected pre-2012, and 33 genes were consistently selected post-2012. We used gene functional predictions^42^ to identify their potential metabolic pathways. Of the 33 consistently selected genes post-2012, ten are involved in amino acid metabolism or aminoacylation, and six are involved in nucleic acid synthesis or degradation (**Fig. 5F**).

Previously, the absence of biosynthesis or auxotrophies for amino acids and nucleotides has been highlighted for microorganisms with streamlined genomes^43,44^. In the streamlined *Nanopelagicus*, auxotrophies for various amino acids^15,45^ coupled with an enrichment of transporters for many small organic nitrogen compounds, including amino acids^15,46,47^ and nucleic acid components^15,45– 47^ are common. Moreover, the histidine pathway was found split between two different strains of *Nanopelagicus* growing in a mixed culture^45^. Our observation of consistent selection on amino acid and nucleic acid metabolism suggests that these genes differentiate the post-2012 strains. Additionally, the low phytoplankton biomass (**Fig. 4H**) might indicate lower influx of fixed nitrogen into the system, which could have cascading effects on the processing of organic nitrogen in abundant microorganisms. Therefore, it appears that biosynthesis, use, and reuse of small organic nitrogen compounds are key in the ecology and evolution of these globally abundant lake bacteria.

Freshwater lakes are focal points on the terrestrial landscape, processing an estimated 70% of net terrestrial carbon production^48^. These ecosystems are stressed by both climate change^49^ and invasive species^50^, but whether lakes will become net sources or sinks of carbon is uncertain^51,52^. The coincidence of the 2012 evolutionary shifts in *Nanopelagicaceae* with both a species invasion and environmental extremes implicates anthropogenic drivers. Given the foundational role of bacteria in aquatic food webs^53^ and the global abundance of *Nanopelagicaceae*^13^, its evolution may have wide-ranging impacts on freshwater ecosystems and organic nitrogen compounds may play a central role in freshwater responses to global change.

### A continuum of ecology and evolution

The interface between ecology and evolution is delineated by species boundaries, but in bacteria species definitions are hotly debated^5^. Using a commonly chosen definition for microbial species boundaries, we found interspecific ecological dynamics mirrored intraspecific evolutionary dynamics, with no emergent boundary delineating ecology from evolution. Should interactions like competition and niche differentiation between strains be considered ecology, or does the fact that they were inferred from observations of genomic change place them in the realm of evolution? Should positive selection of organic nitrogen metabolism genes be considered evolution, or are soft selective sweeps simply evidence of ecological shifts between phenotypically distinct strains? Can we differentiate ecological from evolutionary processes when they occur on the same time scales, in response to the same likely environmental drivers, and across unclear species delineations?

Our two-decade TYMEFLIES dataset, its associated 2,855 species-representative MAGs, and decades of NTL-LTER environmental data raise these questions again and again. We identified seasonal and decadal strain dynamics that could be considered alternately ecology or evolution across diverse and abundant phyla. Other microbiome studies have similarly identified microdiversity at the strain level as key to understanding microbial change. Strains have displayed distinct environmental preferences in anaerobic digesters^54^, oceans^55–58^, and geysers^59^; and strain-level dynamics have been linked with outcomes such as Cyanobacterial toxicity^60^, preterm birth^61^, human health^62^, and cheese rind aroma^63^. Strains have been described alternately by ecological concepts like metapopulations in the subseafloor^64^ and carrying capacity in the human gut^65^, or by evolutionary concepts like modes of speciation in global lakes^66^. In pitcher plant microbiomes, strains were ecologically distinct when they differed by only 100 SNVs^67^. Among all these microbiome studies, sometimes strain dynamics are framed as ecology^55–57,59,60,63,65,67^ and sometimes as evolution^54,58,61,62,64,66^. However, even in plants and animals speciation is not instantaneous and subspecies population structure creates a blurred line between strains and species^68,69^. Therefore, we propose a shift away from framing eco-evolutionary dynamics around feedbacks between distinct processes^2–4^. To better encompass microbial communities, we should frame change as converging along a continuum of ecology and evolution.

## Methods

### Lake Mendota Samples

Lake Mendota is a eutrophic temperate lake located in Madison, Wisconsin (USA)^70^. Integrated samples were collected from the upper 12 m at a 25 m deep location referred to as the central “deep hole” (43°05’58.2”N 89°24’16.2”W). During the summer stratified months, these 12 m samples span the epilimnion layer. Bacteria were collected on 0.2 µm polyethersulfone filters (Pall Corporation), stored at -80°C, and DNA was extracted by a single person after randomizing sample order in 2018-2019 using FastDNA Spin Kits (MP Biomedicals). A detailed description of the study site, sample collection, and DNA extraction procedures is provided by Rohwer and McMahon^6^.

### Metagenome sequencing and assembly

Samples were sequenced by the US Department of Energy Joint Genome Institute (JGI) using a NovaSeq 6000 with an S4 flow cell. Sample metadata is available in Supplementary Data 1, and raw sequencing data is available from the NCBI Sequence Read Archive under Umbrella Project accession PRJNA1056043. Individual metagenome SRA accession numbers are listed in Supplementary Data 1. Read filtering was performed using standard JGI protocols^71^, which are additionally detailed as metadata paired with each sample through the JGI IMG/M website. Briefly, BBDuk^72^ was used to remove adapters and quality trim reads, and BBMap^72^ was used to identify and remove common contaminants. In our analyses we treated the resulting filtered fastq files as the metagenome reads. Single-sample assemblies were also generated by JGI with their standard protocol^71^ using metaSPAdes^73^. These filtered fastq files and single-sample assemblies are available through the JGI Genome Portal under ITS Proposal ID 504350.

### Obtaining and characterizing genomes

Genomes were binned out of metagenomes using the Texas Advanced Computing Center’s Lonestar6 supercomputer and the Launcher utility (version 3.7)^74^. Metagenomic reads were mapped back to sample assemblies using BBMap (version 38.22)^72^, sorted BAM files were created using SAMtools (version 1.9)^75^, and metagenome-assembled genomes were binned using MetaBAT2 (version 2.12.1)^76^. Metagenomic reads from different samples were cross-mapped back to each assembly. Cross-mapping scales exponentially, so it was performed on assemblies and sample reads broken into approximately 50-sample groups of consecutive sample dates, with samples from the same year grouped together. This resulted in 85,684 genome bins. CheckM2 (version 0.1.3)^7^ was used to asses bin quality, including completeness and contamination estimates, and GTDB-tk (version 2.1.1)^77^ was used to assign GTDB taxonomy (release 207)^78^ to all bins. 30,389 genome bins were at least 50% complete and less than 10% contaminated, and these bins were de-replicated to 96% ANI using dRep (version 3.4.0)^8^. To choose 96% as our ANI cutoff, we ran dRep at ANIs ranging from 90 to 99% and examined the resulting number of de-replicated bins, as well as the number of bins from the same assembly that were combined. We chose 96% ANI because very few (one) of the 30,389 bins were combined into an ANI group with a bin created from the same assembly, and because 96% ANI was generally located right before a sudden increase in the total number of genome groups. Our goal was to separate as many species as possible, while combining strains that were so closely related they would compete for mapped reads. Applying a 96% ANI cutoff with dRep resulted in 2,855 representative genomes, which we treated as species in this study.

To quantify the relative abundance of each species in every sample, we mapped all sample reads against the concatenated 96% ANI reference genomes using BBMap (version 38.22)^72^, created sorted BAM files using SAMtools (version 1.9)^75^, and calculated relative abundance using coverM (version 0.6.1)^79^. With the coverM software, we required a minimum read percent identity of 93, proper pairs only, and excluded 1000 bp from each contig end from the calculation. CoverM calculates relative abundance as the mean coverage divided by the mean coverage across all genomes multiplied by the proportion of reads that mapped to the genome, thus normalizing by recovered genome size to estimate the fraction of cells that belong to a given species in each sample. A table of representative MAGs along with taxonomy annotations, quality statistics, and abundance statistics is available as Supplementary Data 2.

To further characterize the genomes, we ran inStrain (version 1.7.1)^80^ using a minimum read ANI of 93%, as recommended by the inStrain documentation given our previous choice of 96% ANI to dereplicate genomes. This software called SNVs and calculated nucleotide diversity, among other metrics. To identify genes we ran prodigal (version 2.6.3)^81^ on each genome separately. We then used Kofamscan (version 1.3.0)^82^ to assign gene annotations from the Kyoto Encyclopedia of Genes and Genomes (KEGG) database (release 107.1)^42^. Additional custom analyses were performed using the R programming language (version 4.1.2)^83^, and relied extensively on the data.table R package (version 1.14.8)^84^, the lubridate R package (version 1.9.3)^85^, and GNU parallel (version ‘Chandrayaan’)^86^.

### Classifying seasonal and long-term change

To classify each species’ abundance pattern as seasonal or not, we started with relative abundances as calculated by coverM (version 0.6.1)^79^ and further corrected any abundance to zero if the genome’s coverage breadth was 70% or less than its expected breadth, as calculated by inStrain (version 1.7.1)^80^. We then applied a centered log ratio transformation to the relative abundance values using the compositions R package (version 2.0-6)^87^. After taking a daily linear interpolation to obtain evenly spaced samples, we detrended the temporal profiles with a cubic fit. Finally, we performed a periodogram analysis by computing the magnitude of the fast Fourier transform. If a peak occurred within 30 days of 365 days we considered it an annual oscillation, and if any of the top five peaks corresponded to an annual period, we classified the species as having a seasonal abundance pattern. We applied this analysis only to the 1,474 species that occurred on least 30 dates over at least 10 years. To classify each species’ nucleotide diversity pattern as seasonal or not, we similarly performed a fast Fourier transform on its inStrain-calculated nucleotide diversity over time. We used the same periodogram analysis to classify it as having seasonal nucleotide diversity or not, and we applied this analysis to the same subset of 1,474 species.

To characterize blooms as more diverse or less diverse, we calculated the Pearson correlation between centered log ratio-transformed relative abundance and nucleotide diversity for the 365 species that had both seasonal abundance and seasonal nucleotide diversity annual oscillations. We considered it a positive correlation (more diverse blooms) if the Pearson correlation was at least 0.35 and a negative correlation (less diverse blooms) if the Pearson correlation was less than or equal to -0.35. We repeated this analysis with up to two weeks of lag and used the highest correlation within that window. We chose 0.35 as a reasonable cutoff after manual examination of the first 150 species’ correlations.

To calculate SNV profiles for each species, we created vectors corresponding to every SNV position in its genome, where the value of each element was the percent of mapped reads that matched the reference genome base at that position in each sample. SNV’s were called using inStrain^80^, and we only applied this analysis to samples where the species’ median coverage was over 10x, as at coverages less than that we observed a drop in the total SNVs called. Therefore, for both long-term and seasonal analysis of SNV profiles, we included only species that had medium coverage over 10x on at least 30 dates over at least 10 years, which resulted in a subset of 263 species. To identify changes in SNV profiles, we created a distance matrix for each species based on Euclidean distances between each sample’s SNV profile using the vegan R package (version 2.6-4)^88^. From this we created a table of time elapsed and Euclidean distance between each sample date.

To identify seasonal patterns in each species’ SNV profiles, we created a daily linear interpolation of pairwise distances between all samples, taking the mean when multiple sample pairs occurred with the same time interval. After detrending with a cubic fit, we performed a periodogram analysis to identify annual oscillations and the presence of seasonal patterns using the same criteria as with our abundance and nucleotide diversity annual oscillation analysis.

To identify long-term change patterns, we subset our pairwise distance table to the distance of each sample from the first sample. We developed a classifier for these temporal profiles of distances between SNV profiles using 11 manually chosen species. We chose our training set to encompass examples of each pattern of change including no change, and to include both high and low numbers of observations. Our classifier criteria was hierarchical: first gradual change was identified, then step change was identified, and finally disturbance/resilience patterns were identified. After training, the classifier was applied to all 263 species above the abundance cutoff. Gradual change was identified if a linear fit to the daily linearly interpolated distances, excluding dates closer than a month to the starting date, resulted in an adjusted R^2^ of at least 0.55. Dates closer than a month to the starting date were excluded because they tended to be highly similar, and a linear interpolation was applied to account for uneven sampling dates, particularly the high frequency of summer sampling in the latter decade of the timeseries. Possible step change locations were identified after excluding dates closer than a month to the starting date and applying an F test to the linearly interpolated distances using the strucchange R package (version 1.5-3)^89^. If a breakpoint was identified by the F test, the means of measured (as opposed to interpolated) before and after distances were different (two-sided Mann-Whitney p-value < 0.01), and the step resulted in a new mean at least 33% higher than the previous mean, a step change pattern was identified. Disturbance/resilience patterns were then identified using outlier distances calculated by the default boxplot statistics in R. If a date’s distance was > 1.5 times the difference between the 3^rd^ and 1^st^ quartile of observed distances a date was considered an outlier, and if outlier values were maintained for at least a month the species was classified as having a disturbance event with resilience.

### Analyzing abrupt change in *Nanopelagicaceae*

To place environmental conditions in 2012 in context, historical environmental data was collected from the North Temperate Lakes Long-Term Ecological Research program (NTL-LTER) through the Environmental Data Initiative (EDI) interface (https://edirepository.org/) and the US Geological Survey (USGS) Water Data for the Nation (https://waterdata.usgs.gov/nwis) using the USGS dataRetrieval R package (version 2.7.14)^90^. EDI datasets analyzed included ice duration^31^; nutrients, pH, and carbon^91^; major ions^92^; water temperatures combined from multiple datasets^93– 97^ as described in Rohwer *et al*.^22^; phytoplankton^98^; and zooplankton^99^ converted to biomass as described in Rohwer, Ladwig, *et al*.^33^. River discharge measurements were obtained from the USGS for the Yahara River, the primary tributary into Lake Mendota (site ID: 05427718)^32^. After exploring all parameters included in these datasets, the occurrence of a hot, dry year with low primary productivity became apparent. Lake heatwaves spanning much of 2012 were confirmed using the 90^th^ percentile definition from Woolway et al.^38^ and the heatwaveR R package (version 0.4.6)^100^.

Relative abundance and nucleotide diversity of the *Nanopelagicus* MAG ME2011-09-21_3300043464_group3_bin69 were calculated as for the seasonal analysis. New SNVs were identified as SNV positions that were called by inStrain^80^ for the first time in a given sample. To identify dates where an unusual number of new SNVs appeared, possibly indicating the emergence of a new strain, the new SNV counts were compared across all sample dates. Initially, high numbers of new SNVs are expected, so outlier dates were identified among the remaining samples after excluding the initial consecutive dates where new SNVs remained in the 4^th^ quantile. Genes under selection were identified using dN/dS and pN/pS ratios as calculated by inStrain^80^. A McDonald-Kreitman test^101^ was used to identify positively selected genes where the bias of unfixed SNVs to be nonsynonymous was lower than the bias of fixed SNVs to be nonsynonymous (pNpS/dNdS < 1), and positive selection was considered statistically significant when the two-sided Fisher p-value was less than or equal to 0.05. A gene was considered consistently selected if it appeared under significant positive selection with high frequency (in the 4^th^ quartile). Consistently selected genes were identified for the pre-2012 and post-2012 time periods separately.

Gene annotations were analyzed in the context of the KEGG pathways^42^ they belonged to. For each potential pathway, all genes present in the genome were visualized with KEGG Pathway Maps (https://www.genome.jp/brite/br08901). When multiple genes that surrounded the selected gene existed in the genome, that pathway was considered a likely annotation. When likely pathways involved amino acid metabolism or aminoacylation, they were considered amino acid-related. When likely pathways involved purine or pyrimidine metabolism, they were considered nucleic acid-related.

## Data Availability

Metagenome sequences are available from the NCBI Sequence Read Archive under Umbrella Project accession PRJNA1056043. Individual metagenome SRA accession numbers are also listed in Supplementary Data 1. The filtered fastq files and single-sample assemblies used in this study are available through the JGI Genome Portal under ITS Proposal ID 504350. The 2,855 species-representative MAGs are also under the NCBI Umbrella Project accession PRJNA1056043. Individual NCBI Genome IDs are listed in Supplementary Data 2. Environmental data is publicly available through the Environmental Data Initiative (https://edirepository.org/)^31,91–99^ and the U.S. Geological Survey’s Water Data for the Nation (https://waterdata.usgs.gov/nwis)^32^. Custom scripts used for data processing are available at https://github.com/rrohwer/TYMEFLIES_manuscript and through Zenodo^102^.

## Acknowledgements

Long-term datasets such as TYMEFLIES rely on researchers who contribute a portion of their time and effort to future projects they may not be involved in. This work would not be possible without the generosity of many, including Lake Mendota sampling leads Angela Kent, Tony Yannarell, Ashley Shade, Stuart Jones, Ryan Newton, Georgia Wolfe, Todd Miller, Emily Kara Read, Lucas Beversdorf, James Mutschler, and the original Microbial Observatory lead Eric W. Triplett. We thank Sarah Stevens for her early input into the ideas pursued here, Peter Golightly for advice on genes under selection data, Tyler Butts for advice on environmental data, and William Ratcliff and Vincent Denef for advice on framing.

## Funding

E. Michael and Winona Foster WARF Wisconsin Idea Fellowship (RRR)

U.S. National Science Foundation (DBI-2011002) (RRR)

The Texas Advanced Computing Center (TACC) at The University of Texas at Austin provided HPC resources that contributed to the research results reported within this paper (http://www.tacc.utexas.edu) (RRR)

U.S. National Institutes of Health (R01-GM116853) (MKirk)

U.S. National Science Foundation (DEB-1831730) (MKirk)

The work (proposal: https://doi.org/10.46936/10.25585/60001198) conducted by the U.S. Department of Energy Joint Genome Institute (https://ror.org/04xm1d337), a DOE Office of Science User Facility, is supported by the Office of Science of the U.S. Department of Energy operated under Contract No. DE-AC02-05CH11231 (MKell)

U.S. Department of Energy Joint Genome Institute (CSP 504350) (KDM)

U.S. Department of Agriculture (WIS01516 and WIS01789) (KDM)

U.S. National Science Foundation (DEB-0702395, DEB-1344254) (KDM)

U.S. National Science Foundation North Temperate Lakes Long-Term Ecological Research program (DEB-9632853, DEB-0217533, DEB-0822700, DEB-1440297, DEB-2025982) (KDM)

U.S. National Science Foundation Microbial Observatory program (MCB-9977903, DEB-0702395) (KDM)

Simons Foundation Investigator in Aquatic Microbial Ecology Award (LI-SIAME-00002001) (BJB)

## Author contributions

R.R.R. and K.D.M. conceptualized the research and obtained initial funding. K.D.M. and B.J.B. provided resources. R.R.R. conducted field and laboratory work and curated data. RRR performed analyses and created visualizations. M.Kirk. advised statistical approaches. S.L.G., M.Kell., K.D.M., and B.J.B. advised analysis approaches. R.R.R. wrote the first draft, and R.R.R. and B.J.B. wrote the final draft incorporating edits provided by M.Kirk., S.L.G., M.Kell., and K.D.M.

## Supplementary Information

**Supplementary Data 1. TYMEFLIES metagenome metadata**. Includes metadata for metagenome samples including JGI, GOLD, and NCBI sample identifiers as well as McMahon Lab identifiers that pair metagenome samples with previous 16S rRNA gene sequencing^6^.

**Supplementary Data 2. TYMEFLIES MAG metadata**. NCBI identifiers corresponding to each species-representative genome, as well as genome quality calculated by CheckM2^7^, taxonomy assigned by GTDB-tk^77^, and average relative abundance calculated by coverM^79^.

**Supplementary Data 3. Consistently selected gene annotations**. KEGG annotations of consistently positively selected genes in a *Nanopelagicus* species that experienced a step change in strain composition in 2012 (ME2011-09-21_3300043464_group3_bin69). Table row order matches heatmap row order in **Fig. 5F**.

